# Polymer brush bilayer under stationary shear motion at linear response regime: A theoretical approach

**DOI:** 10.1101/565796

**Authors:** Mike John Edwards

## Abstract

Statistical mechanics is employed to tackle the problem of polymer brush bilayers under stationary shear motion. The article addresses, solely, the linear response regime in which the polymer brush bilayers behave very much similar to the Newtonian fluids. My approach to this long-standing problem split drastically from the work already published Kreer, T., *Soft Matter*, **12**, 3479 (2016). It has been thought for many decades that the interpenetration between the brushes is source of the friction between the brush covered surfaces sliding over each other. Whiles, in the present article I strongly reject that idea. Instead, here, I show that structure of the whole system is responsible for friction between brush covered surfaces and the interpenetration is absolutely insignificant. Two simple reasons for that are the presence of ambient solvent and also flexibility of the chains. The results of this research would blow one’s mind about how the polymer brush bilayers respond at small shear rates.

## Introduction

Polymers are linear macromolecular structures that are composed of repeating building blocks of atomic or molecular size i.e. monomers.^1,5,19^ Typically, polymers bear 10^4^ to 10^5^ monomers per chain (degree of polymerization) in biological systems.^1,5,19^ In principle, the word polymer is devoted to long linear macromolecules with many monomers. The connectivity between monomers in chain is established through the Covalent bond in which monomers share their valence electron together. In 1827, R. Brown discovered that particles at molecular and atomic scale undergo thermal fluctuations. Since, monomers undergo thermal agitation as a result of collision with solvent molecules, therefore the whole polymer chain undergoes all possible conformations in the course of time. However, by measuring the chain extension in a long time scale, at thermal equilibrium the chain extension reaches a certain average length. This length is determined by *entropic elasticity* (which shrinks the chain) from one hand and the *steric repulsion* among monomers (which stretches the chain) on the other hand.^1,5,19^

One of the most interesting polymeric structures that exists in many biological systems is the polymer brushes.^2^ Polymer brushes are composed of linear chains that are chemically and densely grafted into a surface. The steric repulsion among monomers of nearby chains stretches the chains strongly in the perpendicular direction. The brush like structures could be found as Glycol on the outside of cell membranes as well as aggregan in synovial fluids of mammalian joints. Since thickness of the brush layer is easily controllable by varying molecular parameters, therefore one could tune surface properties by using brushes. Polymer brushes have been investigated theoretically^7,9–11^ and by experiments.^8^ Polymer brush bilayers (PBB) are observed in synivial joints of mammals and it’s said that they are responsible for reduction of friction between the bones sliding over each other. Moreover, it has been also said that the PBBs are responsible for suppression of the mechanical instabilities in synovial joints. To investigate such a system, one would normally take two opposing surfaces with a certain distance and each surface covered by a brush that are similar in molecular parameters. The brushes got to be at intermediately compressed state to make balance between compression and interpenetration. The most simplified motion that resembles the synovial joints motion, could be the shear motion where surfaces move in opposite directions with the same speed. Based on the shear rate, response of the PBBs to the shear forces are classified into two regimes. The regime of low shear rates or the linear response regime and the regime of high shear rates in which non-linear response looms. In this article, I focus on the linear response regime where the PBBs behave like the Newtonian fluids. Note that, here I do not address the regime of high shear rates where the nonlinear effects i.e. shear thinning established. It has been taught, for decades, that the interpenetration between the brushes is responsible for producing friction between sliding brush covered surfaces.^4^ Also, there has been an extensive experimental research on usefulness of PBBs in lubrication.^15–17^ Here, I develop a theory that strongly rejects the significance of interpenetration length in producing friction. Instead, my theory suggests that the structure of whole system is responsible for the friction. My theory is based on the density functional theory framework which has been successfully applied to PBBs.^6,18^ In the next section, I describe the theory in details and make comparisons with the existing theoretical approaches to the same problem.

## Theoretical description

### Newtonian fluids

Let us consider a Newtonian fluid which is restricted between two flat plates of surface area *A* and distance *D*. Under the shear motion the flat plates move in opposite directions with speed *v*. In such a situation, the governing equations are the Stokes equations as follows,

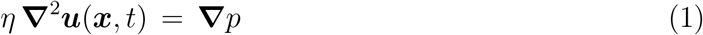

where *η* is the fluid viscosity, ***u***(***x***, *t*) the fluid velocity and *p* the fluid pressure. Assume that the flat plates are in xy plane such that the bottom plate is located at *z* = 0 and the top plate at *z* = *D*. If the shear direction is supposed to be in the x direction, then the following differential equation holds,

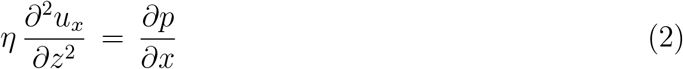

In the absence of external flow, one would set 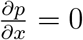. By considering the no-slip boundary conditions i.e. *u*_*x*_(*z* = 0) = *−v* and *u*_*x*_(*z* = *D*) = *v*, the solution of Eq. (2) is,

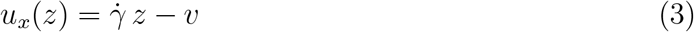

Where 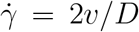 is the shear rate. Therefore, the velocity profile is linear in *z* and vanishes at the center of channel. The stress tensor is given as follows,

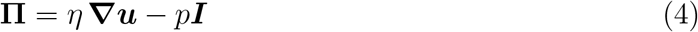

Which implies the following results for normal and shear stresses,

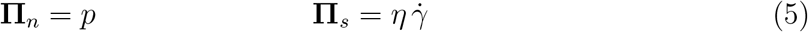

It turns out that, the viscosity and the equation of state of the fluid give a full description of the response under stationary shear motion. That is an important property of the Newtonian fluids. Having reviewed the Newtonian fluids under stationary shear, I will consider PBBs under stationary shear in the following section.

### The PBBs under stationary shear motion

Now let us consider brushes that are grafted onto the flat plates and are suspended in a solvent. The equation of state of a PBB has been calculated^20^ and it is given as follows,

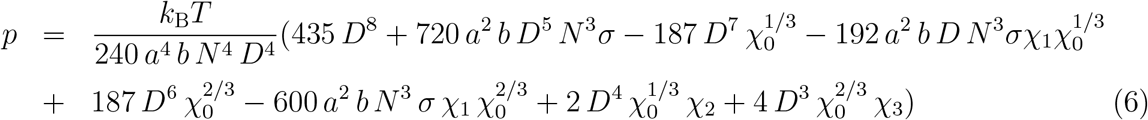

where *a* is the *Kuhn* length or monomer size, *b* the second Virial coefficient, *N* the degree of polymerization, *σ* the grafting density and I have introduced the following volumes for the sake of brevity,

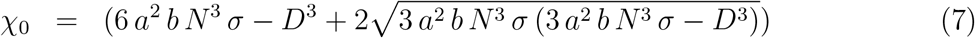

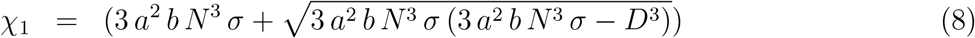

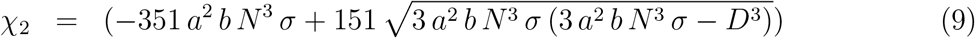

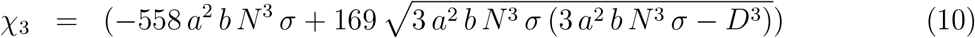

Fig. 1 reveals that the pressure follows the universal power laws as given below,

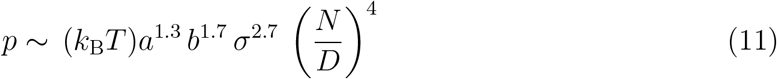

**Figure 1:**
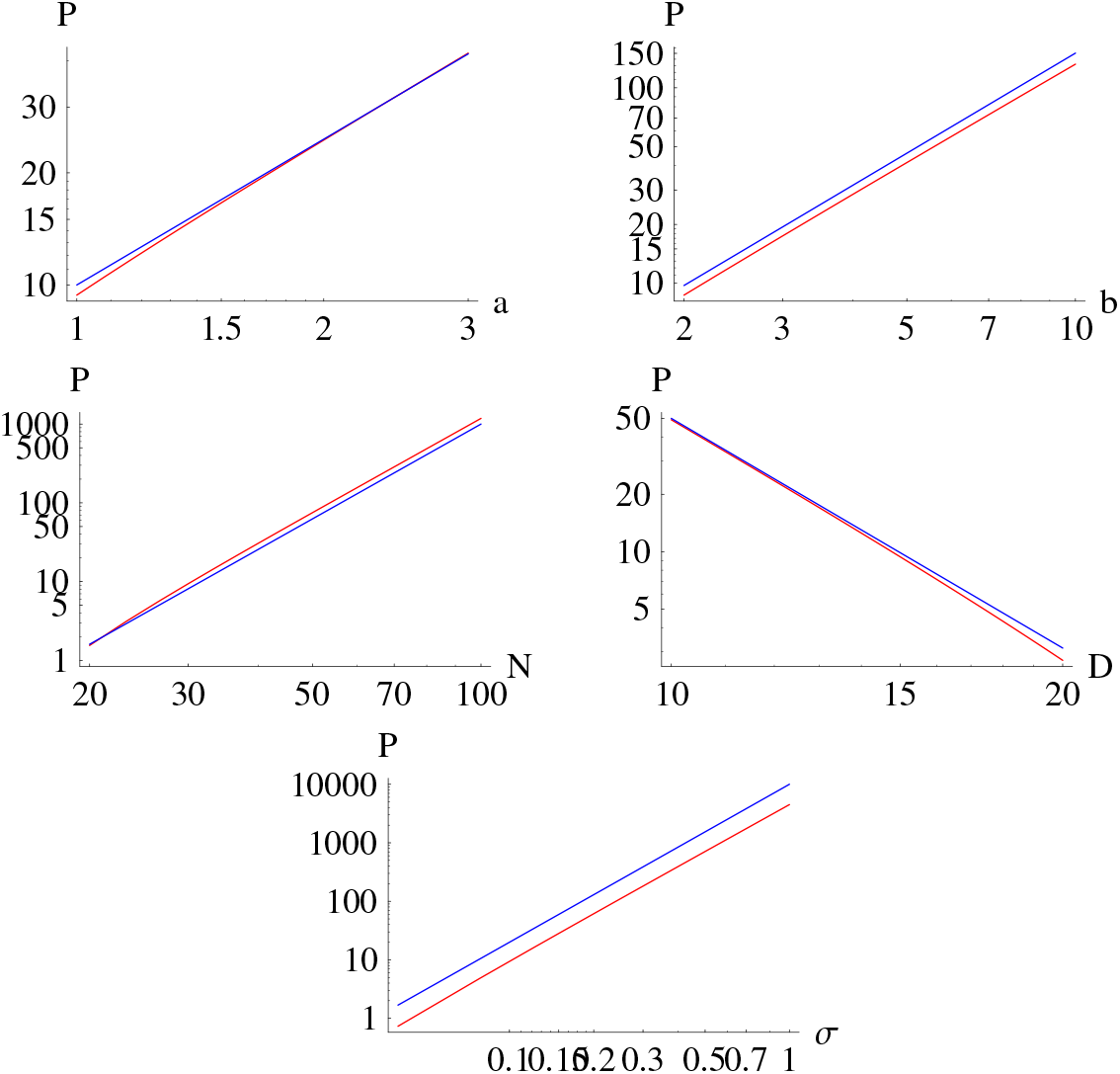
The equation of state of the PBBs in terms of system parameters. The red shows Eq. 6 and the blue shows fitted power law.

The shear forces do not change the equation of state as long as the linear response regime holds. The linear response regime is valid if the Weissenberg number *W* is smaller than unity. The Weissenberg number is a dimensionless number that is defined as 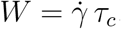, where *τ*_*c*_ = (*N*^2*ν*+1^*a*^2^*ξ/*3*π*^2^ *k*_B_*T*) is the critical time scale below which the chains can relax and *ξ* the monomer friction constant. In principle, *τ*_*c*_ is the largest mode of the Rouse time scale with excluded volume interactions. This is taken into account as the largest relaxation time of polymer chains without hydrodynamic interactions between monomers. ^19^ Therefore, all calculations are valid for (*W* ≪ 1) in which the shear forces are not able to overcome the entropic forces that make monomers diffuse.

So far, I have given a full description of the PBB equation of state as well as the validity range of the linear response regime. According to Eq. (5), we know the normal stress as **Π**_*n*_ = *p*. To calculate the shear stress for the PBBs, one has to get to know about the viscosity of the PBBs. Basically, viscosity is an intrinsic property of any substance. According to the statistical mechanics, the viscosity of any system is equal to its equation of state multiplied by the average time between two successive collisions between its particles i.e *mean free time* or *collision time*.^3^ For PBBs, the collision time of monomers can be calculated as follows. For the PBBs, the mean free path or the average distance in which a monomers moves without collision is 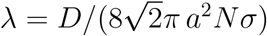 and the diffusion constant of monomers in the Rouse model with excluded volume interactions is (*k*_B_ *T/Nξ*). By dividing the mean free path squared by the diffusion constant, one would get the following time scale as the collision time for monomers in a PBB,

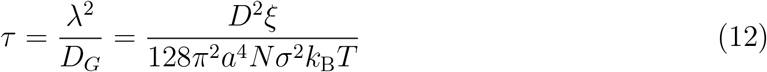

This is the shortest time scale during which monomers are kicked by other monomers. Note here that, the kicks that monomers get from the solvent molecules are considered, and here, I assume that monomers are much larger than solvent molecules. Having calculated the collision time for the PBBs, one would calculate the PBB viscosity as *η* = *pτ*. Eq. (13) shows that the PBBs viscosity follows the universal power laws as given bellow,

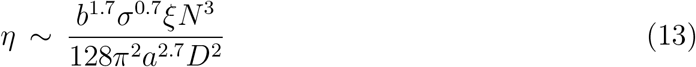

The viscosity increases strongly by *N* and it strongly decreases by *D*. Now that the viscosity as well as the equation of state are calculated, one would calculate the shear stress and the kinetic friction coefficient as following,

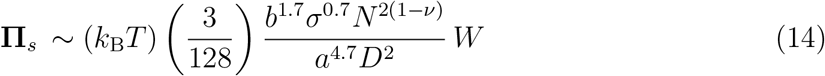

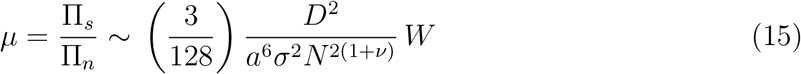

Note here that, in both equations, the Weissenberg number itself scales as *D*^*−*1^. It means that the actual power law of the wall distance is **Π**_*s*_ *∼ D*^*−*3^ and *µ ∼ D*. The chain extensions at low shear are given as follows,^20^

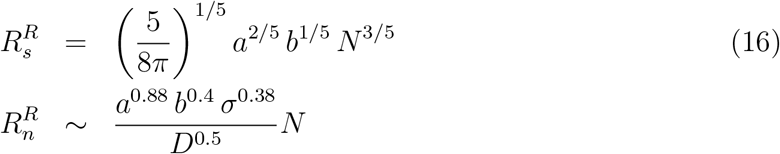

the above relations say that as long as the Weissenberg number does not exceed unity the chain extensions remain the same as equilibrium conditions and they do not become a function of the Weissenberg number.

In the next section, I proceed to make conclusions and discuss the final arguments. Also, I would like to discuss some advantages of my approach to the problem of the PBBs under stationary shear.

## Concluding remarks

In this article, I addressed the polymer brush bilayers under stationary shear motion at linear response regime. At linear response regime the shear forces are weaker than the entropic forces and are not able to change the equation of states. However, the normal and shear stresses still split by presence of the weak shear forces. This is due to the non-isotropic nature of PBBs. This is the linear response regime in which the PBBs behave some what similar to Newtonian fluids. The only thing the PBBs have in common with the Newtonian fluids is the linear dependence of transport quantities upon the shear rate. Clearly, the dependence upon molecular parameters are totally different from those of the Newtonian fluids.

The idea here is that, the whole structure of the PBB system is responsible in producing friction under shear. The complex processes at molecular scale such as collision time and the equation of state are responsible for the friction and definitely the interpenetration is not responsible for that as it is offered in.^4,12–14^ I reject the responsibility of the interpenetration between the brushes in friction and one possible answer could be given as following. It is known that every shear induced forces will be relaxed in the system as long as the linear response regime holds. In these conditions, how to admit that the stress produced in the narrow interpenetration region will be transferred to the walls without being relaxed. Another important argument for rejecting the significance of the interpenetration lays in the nonlinear response regime of the PBBs. In certain conditions, it has been observed (which is discussed in^21^) that at Weissenberg numbers higher than unity, the shear stress follows sublinear (or even super-linear) regime while the interpenetration length completely disappear. However, the shear stress still is non-zero even increases by the shear rate. If the interpenetration was responsible for friction, the shear stress should drop. So it seems irrational to relate these two effects to each other. The viscosity increases strongly by *N*^3^ and weakly by *σ*^0.7^. This reveals that the connectivity among monomers contributes more to viscosity rather than number of chains. The kinetic friction coefficient increases by *D* and decreases by *σ*^*−*2^. Also, it strongly decreases by *N* ^*−*2(1+*ν*)^. These show that the friction decreases by increasing monomers in either ways. Also, it says that increasing the wall distance makes more friction. The problem of PBBs under stationary shear at nonlinear response regime would be the next step of my research. Moreover, the non-equilibrium conditions like shear inversions or Polyelectrolyte brush bilayers could be candidates for the future studies.

## References

(1) Rubinstein, M. and Colby, R.H., Polymer Physics, OUP Oxford (2003)

(2) Advincula, R.C. and Brittain, W.J. and Caster, K.C. and Rühe, J., Polymer Brushes, Wiley-VCH, Weinheim (2004)

(3) Schwabl, F. and Brewer, W.D., Statistical Mechanics, Advanced Texts in Physics, Springer Berlin Heidelberg, 313–316 (2006)

(4) Kreer, T, Polymer-brush lubrication: a review of recent theoretical advances, Soft Matter, 12, 3479 (2016)

(5) Gennes, Pierre Gilles, Scaling Concepts in Polymer Physics, Cornell University Press, 74–75 (1979)

(6) Hirz, S. J., Modeling of Interactions Between Adsorbed Block Copolymers, University of Minnesota, Minneapolis, MN (1988)

(7) Milner, S. T., Polymer Brushes, 251, 4996, 905–914, Science (1991)

(8) Halperin, A. and Tirrell, M. and Lodge, T. P., Tethered chains in polymer microstructures, Macromolecules: Synthesis, Order and Advanced Properties, Springer Berlin Heidelberg, Berlin, Heidelberg, 31–71 (1992)

(9) Szleifer, I. and Carignano, M. A., Wiley-Blackwell, Tethered Polymer Layers, Advances in Chemical Physics, 165–260 (2007)

(10) Milner, S. T. and Witten, T. A. and Cates, M. E., A Parabolic Density Profile for Grafted Polymers, EPL (Europhysics Letters), 5, 5, 413 (1988)

(11) Zhulina, E.B. and Borisov, O.V., Structure and stabilizing properties of grafted polymer layers in a polymer medium, Journal of Colloid and Interface Science, 144, 2, 507–520 (1991)

(12) Galuschko, A. and Spirin, L. and Kreer, T. and Johner, A. and Pastorino, C. and Wittmer, J. and Baschnagel, J., Frictional Forces between Strongly Compressed, Nonentangled Polymer Brushes: Molecular Dynamics Simulations and Scaling Theory, Langmuir, 26, 9, 6418–6429 (2010)

(13) Kreer, Torsten and Binder, Kurt and Müser, Martin H., Friction between Polymer Brushes in Good Solvent Conditions:? Steady-State Sliding versus Transient Behavior, Langmuir, 19, 18, 7551–7559 (2003)

(14) Kreer, T. and Müser, M. H. and Binder, K. and Klein, J., Frictional Drag Mechanisms between Polymer-Bearing Surfaces, Langmuir, 17, 25, 7804–7813 (2001)

(15) Klein, Jacob and Perahia, Dvora and Warburg, Sharon, Forces between polymer-bearing surfaces undergoing shear, Nature, 352, 143 (1991)

(16) Taunton, Hillary J. and Toprakcioglu, Chris and Fetters, Lewis J. and Klein, Jacob, Forces between surfaces bearing terminally anchored polymer chains in good solvents, Nature, 332, 712 (1988)

(17) Klein, Jacob and Kumacheva, Eugenia and Mahalu, Diana and Perahia, Dvora and Fetters, Lewis J., Reduction of frictional forces between solid surfaces bearing polymer brushes, Nature, 370, 634 (1988)

(18) Safran, S., Statistical Thermodynamics Of Surfaces, Interfaces, And Membranes, Addison-Wesley Publishing Company, The advanced book program (1994), p. 229–231

(19) Doi, M. and Edwards, S. F., The theory of polymer dynamics, Clarendon Press. Oxford (1986), p. 95

(20) Edwards, M. J., Polymer brush bilayers at thermal equilibrium: A theoretical study, 2018, https://doi.org/10.1101/316141

(21) Edwards, M. J., Polymer brush bilayer under steady state shear: Density functional theory, scaling theory and molecular dynamic simulation (2022) https://doi.org/10.1101/2022.08.16.504146

